# AP-1 regulates slow-cycling cells, cellular dormancy and chemoresistance in TNBC

**DOI:** 10.1101/2023.11.22.566980

**Authors:** Yang Dong, Jin Bai, Anmbreen Jamroze, Rong Fu, Huilan Su, Wenwen Xia, Shan Wu, Ruifang Liu, Dean G. Tang, Jianjun Zhou

## Abstract

**Background:** Dormant or slow cycling cells (SSCs) pre-exist in tumor and responsible for chemo-resistant and tumor recurrence. Due to their low differentiation and dormancy characteristics, SCCs are resistant to standard chemotherapy and targeted therapy. Label-retaining is a common method used to identify and isolate live SCCs. However, it remains unclear whether different label-retaining methods yield distinct SCC subpopulations. In this study, we investigated that various label-retaining methods result in overlapping yet heterogeneous subpopulations of SCCs. Additionally, we explored the molecular mechanisms regulating dormancy in triple-negative breast cancer (TNBC).

**Methods:** We employed multiple label-retaining methods to simultaneously label MDA-MB-231 cells, thereby generating distinct subpopulations of SCCs. We subsequently analyzed these subpopulations for heterogeneity in cell cycle distribution, drug resistance, invasive capacity, and other characteristics using real-time PCR, flow cytometry, and Transwell assays. RNA-seq analysis was performed to characterize the gene expression profiles of the SCCs. Furthermore, we used real-time PCR, Western blotting, immunofluorescence, and luciferase assays to investigate the role of characteristic AP-1 expression in dormancy regulation. Finally, the therapeutic effects of targeting AP-1 in the treatment of TNBC were assessed using a cell-derived xenograft model.

**Results:** We labeled and separated three overlapping but non-identical SCCs subpopulations. We found that all three SCCs subgroups are cell cycle arrested. Additionally, Violet enriched SCCs showed stronger drug resistance and more G1 phase arrest, while Claret enriched SCCs demonstrated enhanced migratory and invasive abilities, along with more G2/M phase arrest. Furthermore, we observed upregulation of AP-1 expression in SCCs, and the JunB subunit of AP-1 promoted the expression of CDKN1A and GADD45A, thereby maintaining cell cycle arrest. CC-930 can inhibit AP-1 transcriptional activity by suppressing JNK activity, ultimately improving the therapeutic efficacy and prognosis of TNBC when used in combination with chemotherapy drugs.

**Conclusions:** We obtained three subpopulations of SCCs with heterogeneous drug resistance. Our findings suggest that AP-1 plays a regulatory role in dormancy regulation in TNBC, and elucidated the molecular function of JunB subunit. Targeting AP-1 with CC-930 has the potential to improve the treatment and prognosis of TNBC.

**Graphical Abstract:** 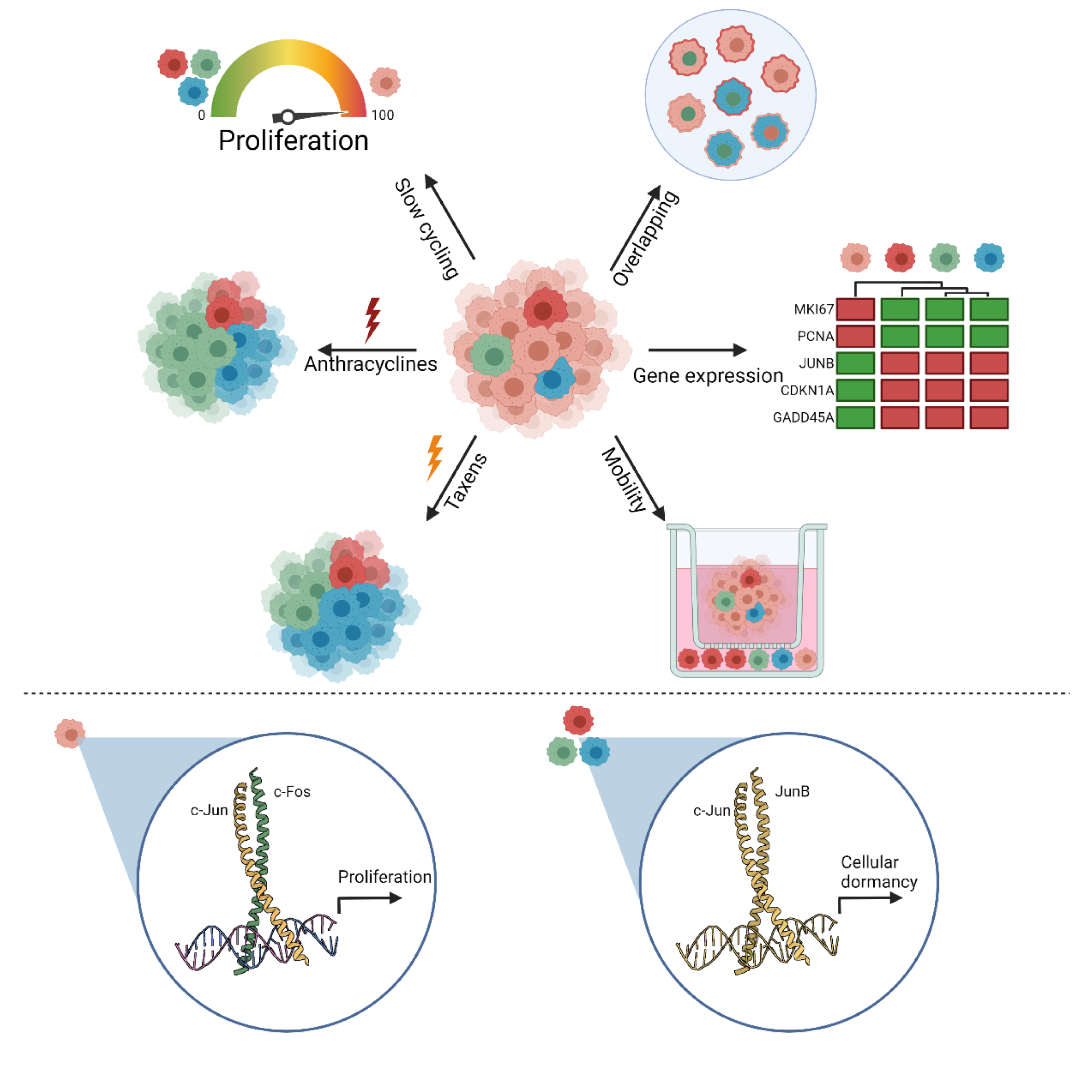

TNBC harbors both fast-cycling cells (FCCs) and functionally overlapping slow-cycling cell (SCC) subpopulations that manifest differential drug sensitivities and motility (A) but are commonly regulated by the JunB-containing AP1 complex (B).

(A). Slow-cycling (quiescent) TNBC cells in culture (a) identified by different label-retaining approaches phenotypically overlap (b), display differential drug sensitivities (c, d) and motility (e) but share common gene expression profiles (f).

(B). Schematic depicting regulation of proliferation in FCCs by the c-Jun/c-Fos AP1 complex (left) and regulation of cellular dormancy in SCCs by c-Jun/JunB AP1 complex.

## 1 BACKGROUND

Breast cancer stands as the most prevalent malignancy among women, bearing the highest rates of morbidity and mortality[1]. Among its variants, triple-negative breast cancer (TNBC) presents a challenge in therapy due to the absence of specific markers, resulting in the poorest prognosis and heightened susceptibility to recurrence and metastasis[2]. Currently, clinical treatments predominantly revolve around targeted drugs or anti-mitotic agents. Previous investigations have uncovered the presence of dormant or slow-cycling cells (SCCs) in both autopsy specimens of tumor patients and in vitro cultured cell lines[3,4]. Due to their dormant and poorly differentiated characteristics, these cells exhibit resistance to standard chemotherapy and some targeted therapies. Additionally, certain tumor cells can enter a dormant state in response to drug interventions to evade drug-induced cytotoxicity, known as therapy-induced cells[5]. Both types of cells reside in situ within tumors, forming minimal residual disease. Some cells may undergo a period of dormancy lasting 5 to 10 years before relapsing[6]. Some residual tumor cells migrate into the bloodstream or disseminate to distant organs, termed circulating tumor cells (CTCs) and disseminated tumor cells (DTCs). These cells also exhibit dormancy characteristics, remaining latent within tissues for extended periods and ultimately contributing to tumor recurrence and metastasis[7–10]. Therefore, research targeting dormant tumor cells is crucial for developing more effective treatment strategies. Currently, research on dormancy in TNBC predominantly focuses on DTCs[11–19], while studies related to dormant tumor cells in situ or SCCs remain limited.

The initial identification of SCCs was primarily achieved through Ki67 staining or BrdU chasing[20,21]. However, these methods are mostly destructive and fail to yield live cells for subsequent culture or analysis. Label retention, a widely used technique, enables the isolation of live SCCs and has been extensively utilized in the study of dormant cancer cells in leukemia, melanoma, colorectal cancer, prostate cancer, and breast cancer[22–26]. In this method, cells from tumor tissues or cell lines are uniformly labeled with a fluorescent dye and then subjected to a prolonged chase period. As cells undergo mitosis, the fluorescent label theoretically divides equally between the daughter cells. SCCs retain more fluorescent labels after multiple mitotic cycles, earning them the designation of label-retaining cells (LRCs). Various fluorescent dyes or markers can be employed for label retention (Fig. 1A, B). Pece et al. utilized the lipophilic fluorescent dye PKH26 to label and isolate human normal mammary stem cells and breast cancer stem cells, noting higher levels of PKH26-retained cells in poorly differentiated tumors compared to well-differentiated tumors[27]. Ebinger et al. labeled patient-derived xenografts of acute lymphoblastic leukemia (ALL) with the protein covalent binding dye CFSE, identifying a rare, long-term dormant cell population resistant to chemotherapy and possessing stem-like properties[28]. Puig et al. adapted a tetracycline-regulated H2B-eGFP fusion protein to label SCCs in colorectal carcinoma, melanoma, and glioblastoma, and further demonstrated the role of TET2 as a key regulator of SCCs in cell number, survival, and tumor recurrence[29]. While label retention methods have been widely employed in the study of adult stem cells or cancer stem cells, there remains a paucity of reports regarding whether different label retention methods yield similar or distinct populations of SCCs. Furthermore, detailed reports on the cellular dormancy mechanisms in breast cancer, particularly triple-negative breast cancer, are still lacking[25,30].

**Fig 1:**
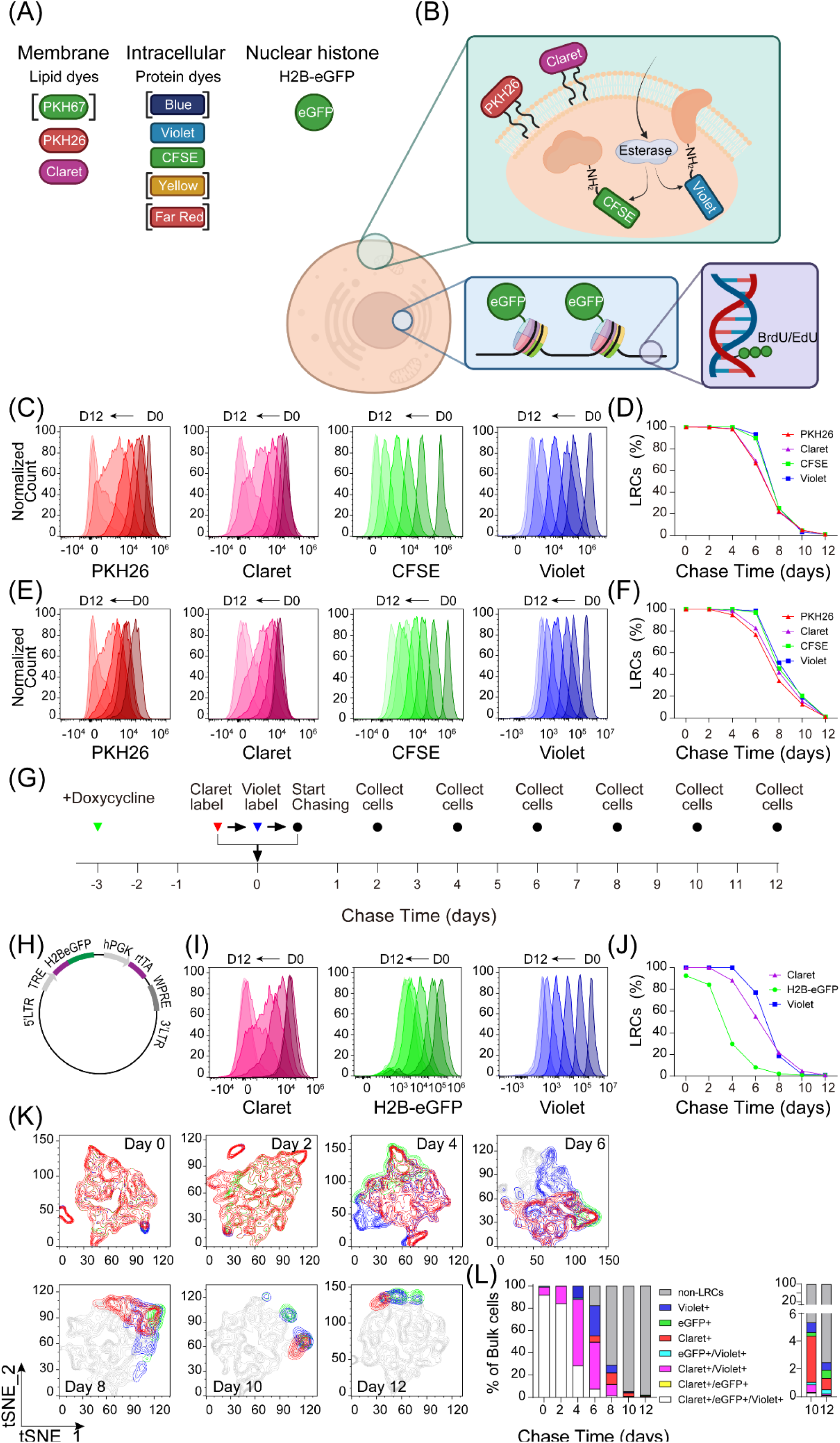
Different LRC strategies identify overlapping but non-identical SCC populations. (A, B) Different labeling methods of SCCs. BrdU/EdU can be incorporated into DNA during S phase instead of thymine. But this method could not label live SCCs for the subsequent experiments. Lipophilic dyes label the phospholipid bilayer of cells through a hydrophobic tail. Protein covalent binding dyes enter the cells by diffusion. The ester group got removed by intracellular esterase, and the fluorescent part of the dye covalently bound to the available amino groups of intracellular protein and intracellular portion of transmembrane protein. eGFP and H2B fusion protein is expressed under the control of tetracycline responding element, thereby traces chromosomes and nucleus. Alternative dyes with different fluorescence are listed on the left. The dyes in parentheses were not used in this study. (C, D) MDA-MB-231 cells and (E, F) SUM159 cells were labeled with PKH26, Claret, CFSE, and Violet separately. Cells were collected and fixed every other day from day 0 to day 12. Fluorescent intensity of each dye was analyzed by flow cytometry. (G) Timeline of tri-labeling strategies. Stable transfected MDA-MB-231 cells were cultured with doxycycline for at least 3 days, then claret labeling and violet labeling were accomplished in order. Cells were collected and fixed every other day from day 0 to day 12. Fluorescent intensity of each dye was analyzed by flow cytometry. (H) Schematics of Tet-On H2B-eGFP expression vector. (I, J) Label retaining effect of Claret, H2B-eGFP and Violet in tri-labeled cells. MDA-MB-231 cells were labeled following aforementioned timeline. Fluorescent intensity of each dye was analyzed by flow cytometry. (K) tSNE plot and (L) stacked histogram of each time point.

In this study, we employed three distinct label retention methods to simultaneously label MDA-MB-231 cells. Following an extended chase period, we observed overlapping yet non-identical enrichment of slow-cycling cells with different fluorescent labels. We examined the similarities and differences among the three subpopulations of slow-cycling cells from various perspectives, including cell cycle, detoxification, and drug resistance. Subsequently, we conducted RNA-seq analysis to elucidate the source of heterogeneity among the three subpopulations of slow-cycling cells. Our findings revealed that all three subpopulations of slow-cycling cells exhibited overexpression of AP-1-related genes, with this heightened expression correlating with the poor prognosis of TNBC. Through molecular investigations, we identified that JunB overexpression could directly activate the expression of genes such as *CDKN1A* and *GADD45A*, inducing cell cycle arrest and cellular dormancy. Finally, we demonstrated that CC-930, an upstream inhibitor of AP-1, could be combined with chemotherapy drugs to achieve enhanced therapeutic effects.

## 2 METHODS AND MATERIALS

### 2.1 Cell lines and labeling

MDA-MB-231 and SUM159 cells were generous gifts from Dr. Zuoren Yu and. HEK293T cell line was a generous gift from Dr. Ke Wei. All cell lines were identified by STR. MDA-MB-231 cells and HEK293T cells were cultured with high glucose DMEM medium supplied with 10% FBS, SUM159 cells were cultured with RPMI-1640 medium supplied with 10% FBS. For PKH26 (Sigma, #MIDI26, USA) and Claret (Sigma, #MIDCLARET, USA) chasing, cells were washed with PBS 3 times, resuspend with Diluent C solution provided with the kit, and incubated with 10 mM PKH26 dye for 5 minutes at room temperature, protecting from light. Afterwards, cells were washed with Diluent C solution 3 times, and cultured in complete medium. For CFSE (Invitrogen, #C34554, USA) and Violet (Invitrogen, #C34557, USA) chase, cells were washed with PBS once, resuspend with PBS and incubated with 5 mM CellTrace dye for 20 minutes at 37 ℃, protecting from light. Afterwards, cells were washed with complete medium twice, then cultured in complete medium. For doxycycline (DOX) chase, cells were treated with 5 mg/ml DOX for at least 3 days in advance, then cultured in DOX-free complete medium. For cells labeled with multiple dyes, perform PKH dyes labeling prior to CellTrace dyes labeling, and culture the labeled cells in DOX-free complete medium (Fig. 1G).

### 2.2 Flow cytometry

For fluorescence activated cell sorting (FACS), cells were labeled and chased as previous described. At the end of chasing, cells were collected and DAPI or PI was used to exclude dead cells during sorting. Cells with top 1% fluorescence intensity were sorted as SCCs, cells with moderate fluorescence intensity were sorted as rapid cycling cells (RCCs), and cells without gating on any fluorescence intensity were sorted as bulk control. 1 * 10^5^ cells of each subpopulation were sorted and used in subsequent experiments. A Beckman Moflo system was utilized to perform FACS.

For flow cytometry analysis, cells were labeled and chased as previous described. PI staining were operated with PI/RNAse staining solution (Invitrogen, #F10797, USA). ALDH activity of each subpopulation was detected by ALDEFLUOR™ Kit (Stemcell, #01700, Canada), and DEAB was used as control. The proportions of senescent cells in each subpopulation were measure by CellEvent™ Senescence Green (Invitrogen, #C10841, USA), with cells treated with 1 µM Etoposide as positive control. A Beckman cytoflex system was used to collect the data, and Flow-Jo v10 was used to analysis the data. The t-SNE module was provided by Flow-Jo v10.

### 2.3 Reverse transcription and quantitative PCR

Total RNA was extracted using Trizol (Invitrogen, #15596018CN, USA) and used to synthesize cDNA using RT reagent Kit with gDNA Eraser, and oligo dT and random hexamer primers were used together (Takara, RR047A, Japan). A Q6 qPCR System was used with TB Green Premix Ex Taq (Takara, RR420A, Japan), and specific pairs of primers (Supplementary Table S1) to detect the indicated transcripts. Relative gene expression was determined by the comparative Ct method.

### 2.4 Immunofluorescence

Tet-on H2BeGFP stabled transfected cells were cultured with DOX for 3 days, chased in DOX-free medium for 12 days, and fixed with 4% PFA. After fixation, cells were permeabilized with 0.1% Triton X-100 in PBS, followed by 2 washed with PBS and blocked with 4% FBS. Primary antibodies (Supplementary Table S2) were incubated with recommended dilution. Secondary antibody and Hoechst 33342 were stained together. Images were collected with ZEISS AX10 system.

### 2.5 Western Blot

Cells were pelleted and washed once with PBS and resuspended in RIPA lysis buffer supplied with protease inhibitor and phosphatase inhibitor. An ultrasonic device was used to disrupt cells, and lysates were centrifuged to collect supernatants. Protein concentrations were quantified with BCA, and 100μg of each component was loaded. Proteins were separated by SDS-PAGE with 5% spacer gel and 12% separating gel, and later transferred to a PVDF membrane. The membrane was blocked with blocking buffer (LI-COR), and primary antibodies (Supplementary Table S2) were incubated with recommended dilution. IRDye-conjugated secondary antibodies were used, and images were collected with an Odyssey system. All the washes were performed with TBST.

### 2.6 Transwell assay

Transwell permeable inserts (Corning) were used for migration and invasion analysis. Tri-labeled cells were chased for 12 days, subpopulation of LRCs and ungated bulk cells were sorted by FACS as described above. Cells were washed and resuspended with 200 μl FBS-free DMEM medium and seeded in the upper chamber of inserts with 2.5 * 10^4^ cells in each well. The inserts used for invasion analysis were coated with 20μg Matrigel (Corning) in advance. 500 ml complete medium was added to each well of 24-well plate as lower compartment, and the inserts were seated soon after. Continued to culture for 24 hours with migration assay, and 48 hours with invasion assay. The inserts were fixed with 1% crystal violet solution for 30 minutes and washed twice with ultrapure water. Cells were counted under microscope with 5 sights for each insert. Images were collected with ZEISS AX10 system.

### 2.7 In vitro chemoresistance

For in vitro chemoresistance assay, tri-labeled cells were chased for 9 days, and 1μM of each drug (TargetMol.USA) were added. Cells were incubated for 72 hours before fixation and analysis with flow cytometry. For anti-AP1 and TAC combination assays, tri-labeled cells were chased for 9 days prior to drug treatment. A combination of 1.867μM docetaxel, 1.724μM doxorubicin, and 35.829μM cyclophosphamide (TargetMol.USA) was named as TAC. 10μM CC-930 or T-5224 (TargetMol.USA) were used solo or in combination with TAC. Cells were incubated for 72 hours before fixation and analysis with flow cytometry.

### 2.8 Plasmid construction

TetOn-H2BeGFP plasmid (pSIN-TRE-H2BeGFP-hPGK-rtTA2) and fastFUCCI plasmid (pBOB-EF1-FastFUCCI-Puro, Addgene #237968) were acquired from Addgene. pLenti-CMV-MCS-mCherry and pLenti-5’MCS-CMV-mCherry were generous gifts from Dr. Ke Wei. pGL3-basic and pRL-SV40 plasmids were generous gifts from Dr. Yandong Li.

For JunB over ovexpression, Homo sapiens JUNB CDS sequence (NCBI Accession No. NM_002229.3) was synthesized and subcloned into multiple cloning sites of pLenti-CMV-MCS-mCherry for lentivirus production. For luciferase assays, AP-1 response element (6x TGAGTCAG) and a minimal TATA-box promoter were synthesized and cloned into pGL3-basic. This constructed plasmid was named pGL3-AP1re and used to detect AP-1 inhibition activity of CC-903 and T5224. Same synthesized sequence was also cloned into pLenti-5’MCS-CMV-mCherry, replaced whole CMV prompter. Constructed plasmid was named pLenti-AP1re-mCherry and used to trace AP-1 activated cells in vitro.

2 kb upstream of transcription initiation site was considered as promoter region. Promoter region of Homo sapiens CDKN1A (NCBI Gene ID: 1026) and GADD45A (NCBI Gene ID: 1647) were synthesized and cloned into pGL3-basic respectively. Constructed plasmid was named pGL3-pCDKN1A and pGL3-pGADD45A, and used to detect JunB’s transcriptional activation of both genes.

All DNA fractions and siRNA were synthesized by Sangon Biotech, Shanghai. All constructs were confirmed by Sanger sequencing.

### 2.9 Dual luciferase assay

HEK293T cells were transducted with corresponding plasmids as shown in the illustration with Lipofectamine 2000 (Invitrogen, #11668019, USA). After 72 hours, cells were collected and lysed with Passive Lysis Buffer provided with Dual-Luciferase Reporter Assay System (Promega, #E1910, USA). Both Firefly luciferase activity and Renilla luciferase activity were measured with Glo/Max luminometer following the manual. Relative luciferase activity was calculated using Firefly luciferase luminous intensity / Renilla luciferase luminous intensity (Luc/RLuc) and normalized with control group.

### 2.10 Colony formation

500 FACS sorted PKH26^high^ cells or PKH26^low^ cells were plated in 12-well plate and cultured with 500μl complete medium overnight. 1μM CC-930 or equivalent DMSO were added, and half medium were replaced every three days. After 12 days of culture, cells were washed with PBS twice and stained with 1% crystal violet solution for 30 minutes and washed twice with ultrapure water. Images were obtained with Bio-Rad Gel Doc system.

### 2.11 SA-β-GAL staining

FACS sorted cells were plated in 4-well chamber slide and cultured overnight. Cells were washed with PBS once and fixed with 500μl β-Galactosidase fixation buffer (Beyotime, #C0602, China). After fixation, cells were washed with PBS for 3 minutes for 3 times, and stained with 500μl mix working staining solution supplied with the kit at 37 ℃ overnight. Images were collected with ZEISS AX10 system.

### 2.12 Tissue microarray (TMA)

TMA (Cat#: TNBC-1202) with 60 paired TNBC samples and adjacent normal tissues were obtained from Shanghai Outdo Biotech, China. Immunohistochemical staining for JunB was performed according to the commercial protocol (Outdo Biotech, Shanghai, China) using anti-JunB antibody (Sigma #HPA019149, USA). The positive proportion of staining was classified as 0, <5%; 1, 5–25%; 2, 25–50%; 3, 50–75%; 4, 75–100%. The staining intensity was then classified as 0, negative; 1, weak; 2, moderate; 3, strong. The total score of JunB staining was determined through proportion scores multiplied by intensity scores. The research on the use of human tissue samples was approved by the Medical Ethics Committees of Shanghai East Hospital, Tongji University.

### 2.13 RAN-seq

Cell were chased and sorted, and RNA was extracted as described above. A total amount of 1μg RNA per sample was used as input material for the RNA sample preparations. Sequencing libraries were generated using NEBNext® UltraTM RNA Library Prep Kit for Illumina® (NEB, USA) following manufacturer’s recommendations and index codes were added to attribute sequences to each sample. The clustering of the index-coded samples was performed on a cBot Cluster Generation System using TruSeq PE Cluster Kit v3-cBot-HS (Illumia) according to the manufacturer’s instructions. After cluster generation, the library preparations were sequenced on an Illumina Hiseq platform and 125 bp/150 bp paired-end reads were generated.

### 2.14 Survival analysis

Metadata of clinical outcome and gene expression data was obtained from TCGA-BRCA cohort. Patients were divided into two groups based on gene expression levels. Survival analysis was performed using the survival and survminer packages in R. The Kaplan-Meier method was used to estimate survival functions, and the log-rank test was employed to compare survival curves between groups.

### 2.15 Xenograft model

1 × 10^6^ MDA-MB-231 cells were resuspended with 50μl PBS and 50μl Matrigel and injected into the fourth mammary fat pad on the right flank of female nude mice. When the tumor volume reached approximately 300 mm^3^ at approximately 1 week, the mice were randomly assigned into control group, CC-930 group (4 mg/kg), TAC (5 mg/kg/week docetaxel, 3.33 mg/kg/week doxorubicin, and 33.33 mg/kg/week cyclophosphamide) group and CC-930 + TAC combination group. CC-930 was given to mice orally, on a daily basis for 21 days. TAC was given to mice i.p. once a week. The tumor volume and body weight were measured three times a week until two days after the final treatment. Tumor volume was calculated as 0.5 × width^2^ × length. Mice were then sacrificed, and tumors were removed, weighted, and fixed for immunohistochemistry.

### 2.16 Synergy Index Analysis

Experiments were performed using incremental dosing of each drug for four weeks. For combination studies, the tumor volume data was annotated as viability data and were loaded into the free software Synergy Finder Plus[31]. The software generated surface response plots by comparing the tumor volume inhibition data to a drug combination reference model obtained from the effect of each drug alone. Synergy Finder implemented a bootstrapping method to compare viability data. We considered a *p*-value of less than 0.05 to establish statistical differences between drugs combinations.

### 2.17 Statistical analysis

Statistical analyses were performed using GraphPad Prism software or using R. In general, student’s *t*-test, paired or unpaired two-tailed *t*-test were used to calculate the statistical significance between comparisons depending on the data type. A Chi-square test was performed to calculate the statistical significance of abdominal metastasis incidence. *P*<0.05 is considered statistically significant.

## 3 RESULTS

### 3.1 Different label strategy identified overlapping but not identical slow cycling cells

To answer the question whether different label retaining method would result in different LRCs subpopulation, we utilized different fluorescent dyes to label and enrich LRCs. We chose lipophilic dye PKH26 and Claret, and protein affinity dye CFSE and Violet as representatives (Fig. 1A, B). First, MDA-MB-231 cells and SUM159 cells were labeled with these dyes separately. After 12-days of chasing, all four fluorescent dyes were found to be able to label and enrich SCCs (Fig. 1C-F and Supplementary Fig. 1). Next, we labeled MDA-MB-231 cells and SUM159 cell with the four dyes simultaneously and compared SCCs subpopulations with each other. Unsurprisingly, PKH26 label and Claret label had good correlation, even after 12 days of chasing, PKH26 enriched SCCs and Claret enriched SCCs still displayed good correlation. Similar results were also found between CFSE and Violet enriched cells. Nevertheless, the initially observed correlation between cells labeled with disparate dye types diminished rapidly over the course of a 12-day chase, underscoring the existence of heterogeneity among SCCs labeled with diverse dye categories (Supplementary Fig. 2).

Next, we employed additional fluorescent labeling to isolate a third SCC subpopulation. To this effect, Enhanced Green Fluorescent Protein (eGFP) was fused with Histone 2B and driven by the tetracycline response element. Upon the introduction of doxycycline into the culture environment, the H2B-eGFP fusion protein was expressed, thereby labeling the nuclei[29] (Fig. 1G, H). After a long period of chasing, we identified another subpopulation of SCCs. To investigate the heterogeneity of different SCCs subpopulations, we labeled Tet-on stable transfected MDA-MB-231 cells and SUM159 cells with lipophilic dye Claret and protein affinity dye Violet (Fig. 1I, J). Following a 12-days of chasing, we observed greater overlap between H2B-eGFP-enriched SCCs and Violet-enriched SCCs, with Claret-enriched SCCs demonstrating less overlap with the other two subpopulations (Fig. 1K and Supplementary Fig. 3). Nonetheless, there remained a small population of cells that retained Claret, H2B-eGFP, and Violet labels concurrently, even after the 12-day chase (Fig. 1L). This sparse subpopulation suggested a shared origin among different SCC subpopulations.

### 3.2 Common phenotypes of SCCs: cell cycle arrested and enhanced stemness

Upon identifying three distinct subpopulations of SCCs, we sought to compare and contrast their gene expression profiles and phenotypes. We first examined the cell cycle distribution of those cells. qPCR showed that three subpopulation of SCCs shared similar gene expression pattern of cell cycle related genes. Most cyclin dependent kinases, including *CDK1*, *CDK2* and *CDK6* were down regulated. Most cyclins, including *Cyclin E1*, *Cyclin E2*, *Cyclin A2*, *Cyclin B1* were down regulated as well. Conversely, *CDK4* and *Cyclin D1*, two genes that thought to be involved in G1 cycle arrest[32], had a up regulated tendency. Genes in Cip/Kip family, including *CDKN1A*, *CDKN1B*, *CDKN1C* were obviously up regulated, and *CDKN2C* was down regulated (Fig. 2A). The alterations in Cdk inhibitors (CKIs) expression further confirmed that SCCs were undergoing G0/G1 cell cycle arrest[33]. Cell proliferation associated genes *MKI67* and *PCNA* were down regulated in all three subpopulations of SCCs, and immunofluorescence showed that H2B-eGFP enriched SCCs had lower expression of Ki67 and p-H3 (Fig. 2B), indicating that most of these SCCs were arrested in the G1 phase of mitosis.

**Fig 2:**
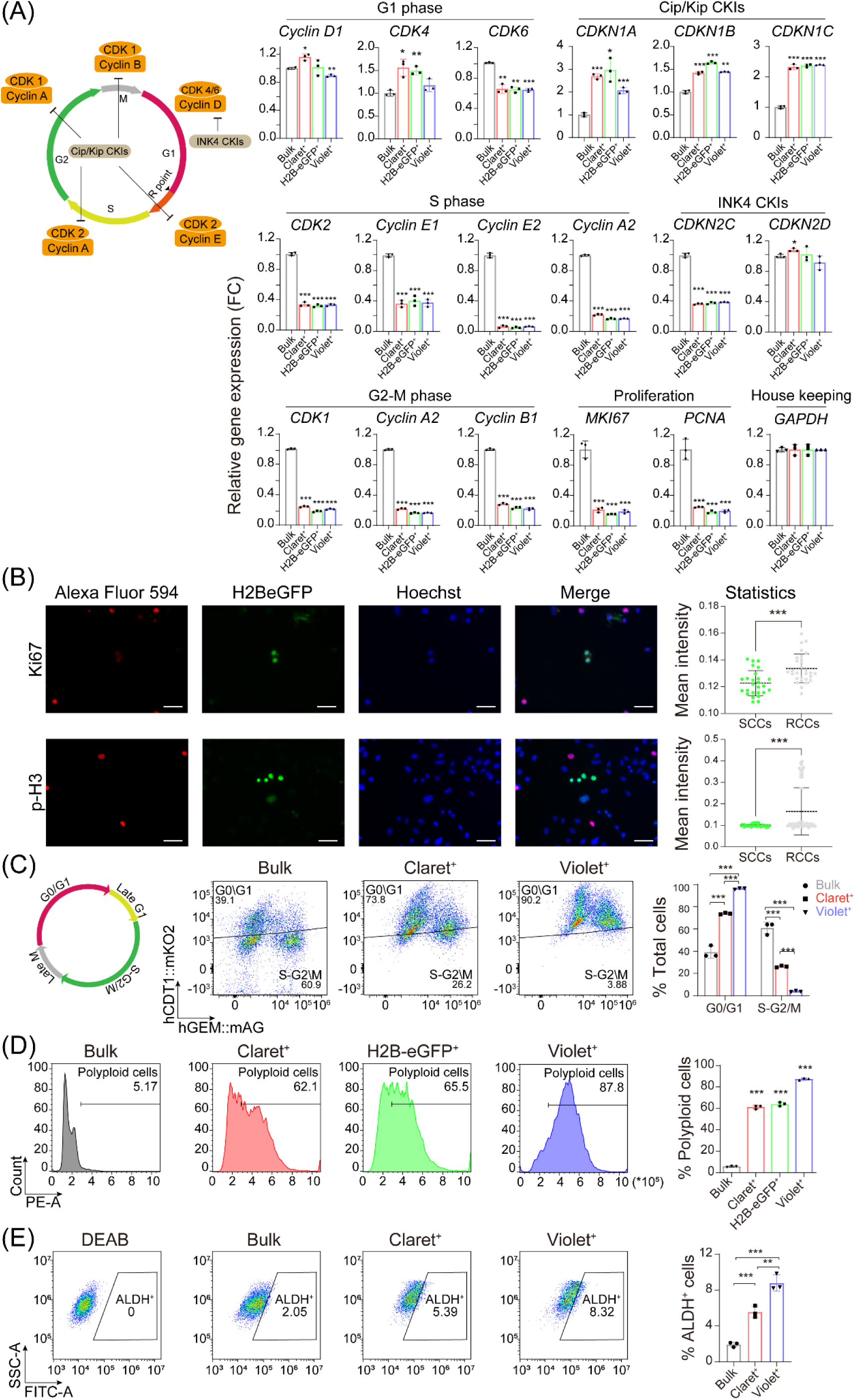
Biological and functional properties of SCCs. (A) 3 SCCs populations showed similar gene expression pattern in cell cycle related genes. Tri-labeled MDA-MB-231 cells were chased for 12 days. Each SCCs population was sorted by FACS, and bulk cells were also sorted as control. Relative gene expressions of SCCs and bulk cells were analyzed by RT-qPCR. (B) Immunofluorescence showed H2B-eGFP enriched SCCs had lower expression of p-H3 and Ki67 after 12 days of chasing. Tet-On H2B-eGFP transfected MDA-MB-231 cells were incubated with doxycycline for 3 days and chasing without doxycycline for 12 days. Cells were fixed and permeabilized, and stained with corresponding antibody. Scale bar = 20 μm. (C) Cell cycle of each population was monitored by FUCCI. FUCCI transfected MDA-MB-231 cells were labeled with Claret and Violet and chased for 12 days. Equal amount of bulk cells, Claret^+^ SCCs and Violet^+^ SCCs were collected and cell cycle distributions were analyzed. Claret^+^ SCCs were spread among different phases, while Violet^+^ SCCs were mostly arrested in late G1 phase. (D) Polyploid cells were more enriched in SCCs compared with bulk MDA-MB-231 cells. Tri-labeled MDA-MB-231 cells were chased for 12 days. Cells were fixed and stained with PI. Equal amount of bulk cells, Claret^+^ SCCs and Violet^+^ SCCs were collected and cells DNA content >4N were considered as polyploid cells. Violet^+^ SCCs enriched more polyploid cells then H2B-eGFP^+^ SCCs and Claret^+^ SCCs. (E) ALDH^+^ cells were more enriched in SCCs compared with bulk MDA-MB-231 cells. MDA-MB-231 cells were labeled with Claret and Violet and chased for 12 days. Equal number of bulk cells, Claret^+^ SCCs and Violet^+^ SCCs were collected and ALDH^+^ cells were identified by DEAB control. Violet^+^ SCCs enriched more ALDH^+^ cells then Claret^+^ SCCs. Data are represented as mean ± SD. *p ≤0.05; **p ≤0.01; ***p ≤0.001, Student’s t test.

We subsequently employed the FastFUCCI system to analyze the cell cycle phase of individual cells. In eukaryotic cells, the licensing factor CDT1 reaches its peak during the G1 phase, subsequently precipitating rapidly in the S phase. Conversely, its inhibitor, Geminin, exhibits heightened expression levels throughout the S and G2 phases, however, it decreases during late mitosis and the G1 phase. These alternating protein expressions are regulated by the alternate activation of E3 ubiquitin ligase SCF^Skp2^ and APC/C^Cdh1^. Through the Fluorescence ubiquitin cell cycle indicator (FUCCI) technology, the fluorescent proteins mKO and mAG are fused to the destabilized sequences of SCF^Skp2^ and APC/C^Cdh1^ respectively, providing an accurate method to track the cell cycle of individual cells[34,35]. FUCCI analysis revealed that a majority of both Claret-enriched SCCs and Violet-enriched SCCs were arrested in the G1 phase (Fig. 2C). However, unlike each other, a mere 10% of Violet-enriched SCCs entered the S phase or G2/M phase, while nearly 30% of Claret-enriched SCCs transitioned into the S phase and G2/M phase. Based on these findings, we can conclude that these three SCC subpopulations were dormant and arrested within the G1 phase.

Furthermore, we examined the association between SCCs and other functional cellular subsets. Firstly, we assessed the detoxification capacity of SCCs using the Aldeflour assay[36]. We found ALDH^+^ cells to be more prevalent in both Claret and Violet enriched SCCs than in bulk cells. Moreover, they were more abundant in Violet enriched SCCs than in Claret enriched SCCs (Fig. 2E). Next using senescence-associated β-galactosidase (SA-β-GAL) staining, we investigated the link between SCCs and senescent cells. Our analysis revealed few senescent cells in SCCs, signifying that the three SCC subpopulations we identified were not senescent cells with irreversible growth arrest (Supplementary Fig. 4A-C). Then using PI staining, we found a large number of abnormal polyploid cells existed in all three subpopulations of SCCs[37], especially in Violet enriched SCCs (Fig. 2D). We extracted genomic DNA from three subpopulations of SCCs and observed that the Violet-enriched SCCs contained a higher amount of genomic DNA compared to the other subsets (Supplementary Table S3). Finally, we utilized a transwell assay to evaluate the migration and invasion abilities of SCCs. Claret enriched SCCs showed enhanced mobility. While H2B-eGFP and Violet enriched SCCs exhibited higher mobility than bulk cells, although the difference was not statistically significant (Supplementary Fig. 4D, E). Collectively, these data suggest that SCCs from the three subpopulations exhibit similar yet divergent phenotypes.

### 3.3 Different subpopulations of SCCs had diverse chemo-resistance

We subsequently investigated the chemoresistance exhibited by three subpopulations of SCCs to a spectrum of chemotherapeutic drugs. Thirteen chemotherapeutic agents were chosen, falling under three primary categories: taxanes, anthracyclines, and DNA damage agents. These encompass the prevalent drug types utilized in the clinical treatment of TNBC[38,39] (Fig. 3A). We observed distinct drug resistance profiles among the three subpopulations of SCCs after 3 days of treatment. Initially, all three subpopulations of SCCs exhibited resistance to taxanes, with Violet-enriched SCCs demonstrating greater resistance compared to the others (Fig. 3B). Similarly, in contrast to the other subpopulations, H2B-eGFP-enriched SCCs exhibited higher resistance towards anthracyclines (Fig. 3C). Claret-enriched SCCs displayed reduced chemoresistance compared to the other two subpopulations, indicating a lesser detoxification ability. Additionally, SCCs did not exhibit significant enrichment after treatment with PARP inhibitors (Olaparib and Veliparib) and ATR inhibitor AZD6738, whereas significant enrichment was observed after treatment with ATM inhibitor CP-466722 (Fig. 3D and Supplementary Fig. 5). qPCR analyses indicated suboptimal function in the ATR pathway across all SCC subpopulations, whereas *ATM* and *CHEK2* appeared upregulated (Fig. 3E).

**Fig 3:**
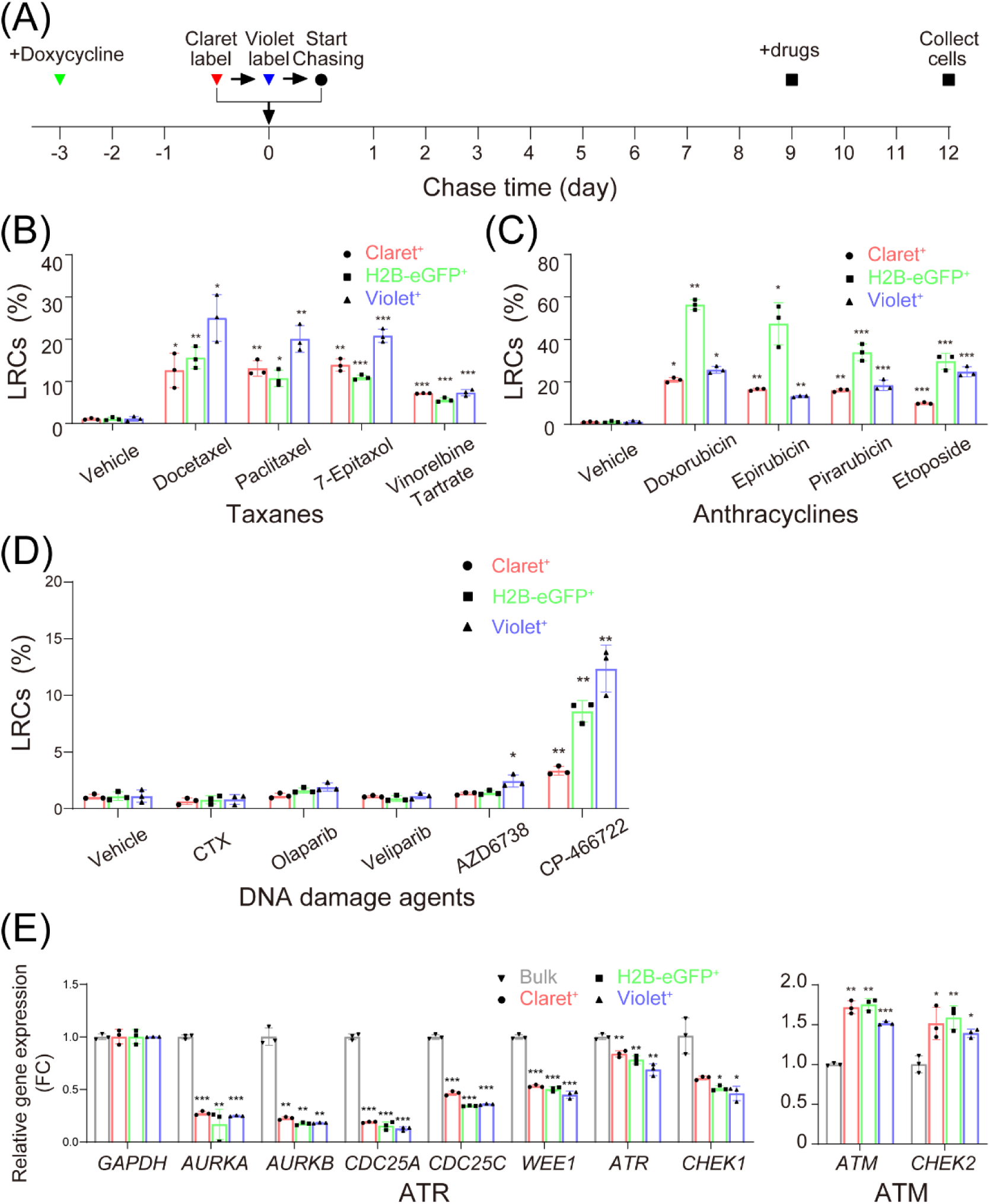
Diverse SCCs have different chemotherapeutic tolerances. (A) Timeline of chemo-resistance assay. Tri-labeled MDA-MB-231 were chased for 9 days, and 1 μM each drug was added respectively. After 3 days of culture cells were collected and analyzed with flow cytometry. (B) More Violet^+^ SCCs retained after taxane treatment, indicating that Violet^+^ SCCs were more resistant to taxanes. (C) More H2B-eGFP ^+^ SCCs retained after anthracycline treatment, indicating that H2B-eGFP ^+^ SCCs were more resistant to anthracyclines. (D) SCCs didn’t show significant retaining after PARP inhibitors treatment. (E) SCCs were up-regulated in *ATM* and *CHEK2*, and down-regulated in ATR pathway. Tri-labeled MDA-MB-231 cells were labeled and chased for 12 days. Each SCCs population was sorted by FACS, and bulk cells were also sorted as control. Relative gene expressions of SCCs and bulk cells were analyzed by RT-qPCR. Data are represented as mean ± SD. *p ≤0.05; **p ≤0.01; ***p ≤0.001, Student’s t test.

### 3.4 Enhanced Correlation in Gene Expression Patterns among Different Subpopulations of SCCs

To delve into the heterogeneity among different subpopulations of SCCs, we conducted a comprehensive bulk RNA-seq using each individual population of SCCs along with ungated bulk cells sorted through FACS (Fig. 4A). This analysis showed that the three subpopulations of SCCs were similar regarding their high Pearson correlation coefficient and shared a majority of differentially expressed genes (DEGs, Supplementary Fig. 6B). The GSVA analysis revealed a significant positive correlation of SCCs with the MAPK signaling pathway and PI3K-AKT signaling pathway, contrasted by the negative correlation with DNA replication, DNA damage repair, cell cycle and cell cycle checkpoints (Fig. 4B). The deficiency in these negatively correlated pathways aligns with a dormant phenotype.

**Fig 4:**
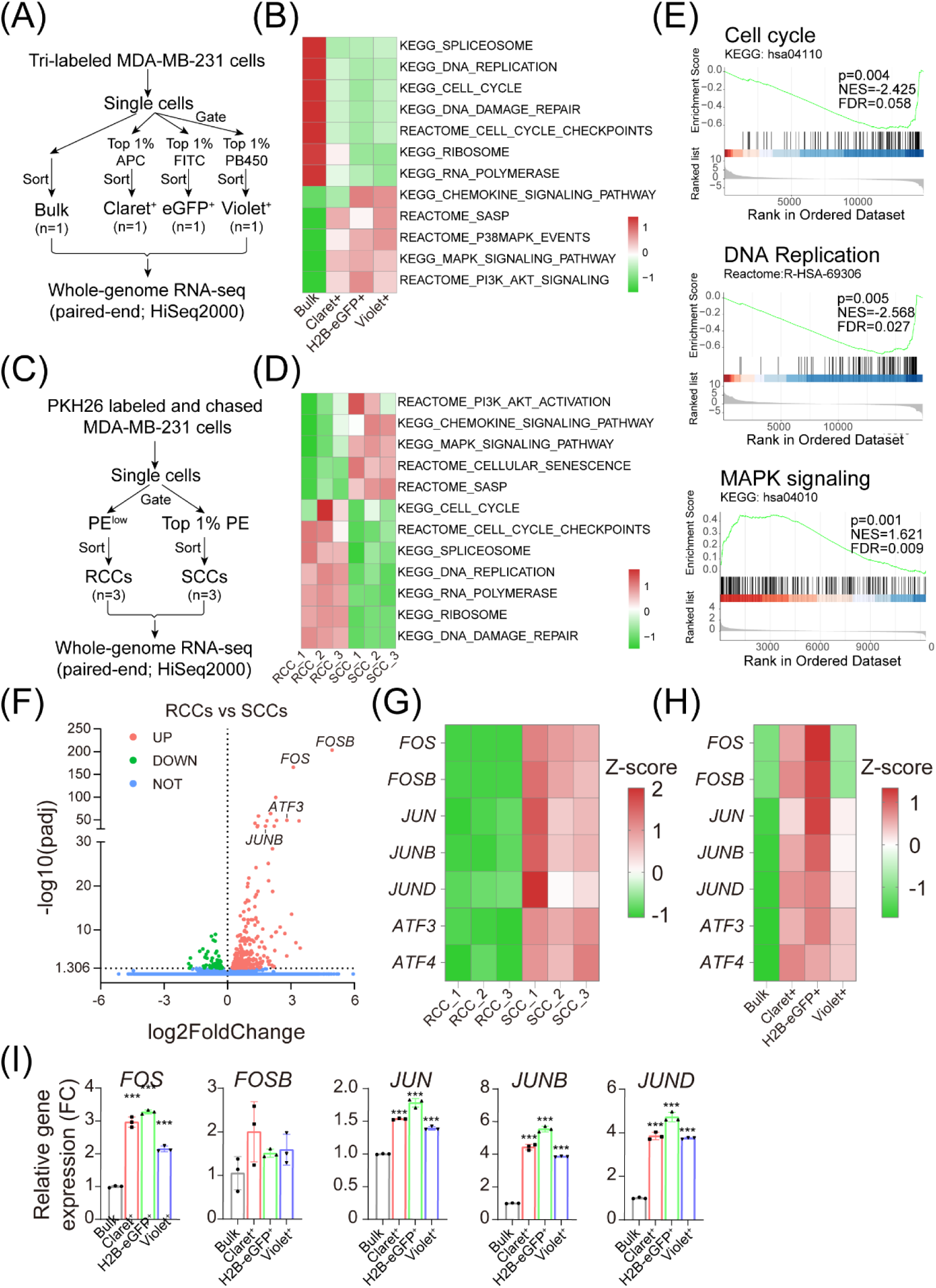
RNA-seq reveal SCCs are enriched in AP-1 signaling and anti-stress mechanisms. (A, C) Flow chart of FACS and RNA-seq. Tri-labeled MDA-MB-231 were chased for 12 days. Top 1% of each population were sorted and names as distinct SCCs subpopulation. Meanwhile, cells that were not gated with any fluorescence channel were sorted as Bulk cell control. Each subpopulation was collected for one sample. MDA-MB-231 cells were labeled and chased with PKH26 for 12 days. Top 1% PE^+^ cells were sorted and named as SCCs, and PE^low^ cells were sorted and names as RCCs. Each subpopulation was collected with triple replications. (B, D) GSVA analysis showed similar pathway enrichment in three SCCs subpopulations. (E) GSEA analysis showed MAPK signaling pathway were up-regulated in PKH26^+^ SCCs, and cell cycle and DNA replication associated pathways were down-regulated in PKH26^+^ SCCs. (F) Volcano plot of DEGs in PKH26^+^ SCCs. FOS and FOSB were the most remarkable up-regulations genes. (G, H) Heatmaps of AP-1 subunits’ expression in RNA-seq data. (I) AP-1 subunits were up-regulated in SCCs. Tri-labeled MDA-MB-231 cells were labeled and chased for 12 days. Each SCCs population was sorted by FACS, and bulk cells were also sorted as control. Relative gene expressions of SCCs and bulk cells were analyzed by RT-qPCR.

We continued our investigation into SCCs’ regulatory mechanisms by performing an additional RNA-seq, this time identifying PKH26+ cells as SCCs and PKH26 low cells as controls for rapid cycling cells (RCCs, Fig. 4C). Our findings showed PKH26+ SCCs shared a similar negative correlation with cell cycle, DNA replication, mRNA splicing and ribosome biogenesis. And also we found positive correlation with MAPK pathway and PI3K-AKT activation as previously observed, along with extracellular matrix organization, chemokine signaling and senescence associated secretory phenotype (Fig. 4D, E and Supplementary Fig. 6C). We also identified the AP-1 subunit *FOS* and *FOSB* as the most upregulated DEGs in PKH26^+^ SCCs (Fig. 4F). Furthermore, other subunits of AP-1, namely *JUN*, *JUNB* and *JUND*, were also found upregulated in SCCs (Fig. 4G, H). We finally validated the expression of AP-1 subunits in three subpopulations of SCCs using qPCR (Fig. 4I), deriving that c-Fos family genes *FOS*, *FOSB* as well as c-Jun family genes *JUN*, *JUNB* and *JUND* were upregulated throughout all three subpopulations. Taking the entirety of the data into account, we hypothesize that AP-1 might be a crucial regulator of TNBC cellular dormancy.

### 3.5 Upregulation of AP-1 subunits in TNBC is a predictor of poor prognosis

To explore the correlation between the expression of AP-1 subunits and the survival of TNBC patients, we categorized the expression levels of AP-1 subunits as high or low using the median gene expression cutoff in the TCGA-BRCA cohort. Initially, we observed that TNBC patients with high expression of FOS or FOSB exhibited worse overall survival (Fig. 5B, C). We further investigated overall breast cancer patients and found no significant difference in overall survival between those with high and low expression of *FOS* or *FOSB*. Considering that ER status is a pivotal factor affecting the survival of BC patients[40], we hypothesized whether the association between high expression of *FOS* or *FOSB* and poor prognosis is related to ER status. Our analysis revealed that high expression of *FOS* or *FOSB* was associated with poor prognosis in ER-negative patients but had no impact on ER-positive patients (Supplementary Fig. 7A). Furthermore, we identified a subset of TNBC patients with concurrent high expression of *FOS* and *FOSB*, who exhibited worse prognosis compared to others (Fig. 4D-F). We validated our findings in the METABRIC-TNBC database (Supplementary Fig. 7B). These results indicate that upregulation of *FOS* or *FOSB* serves as a specific predictor of poor prognosis in TNBC.

**Fig 5:**
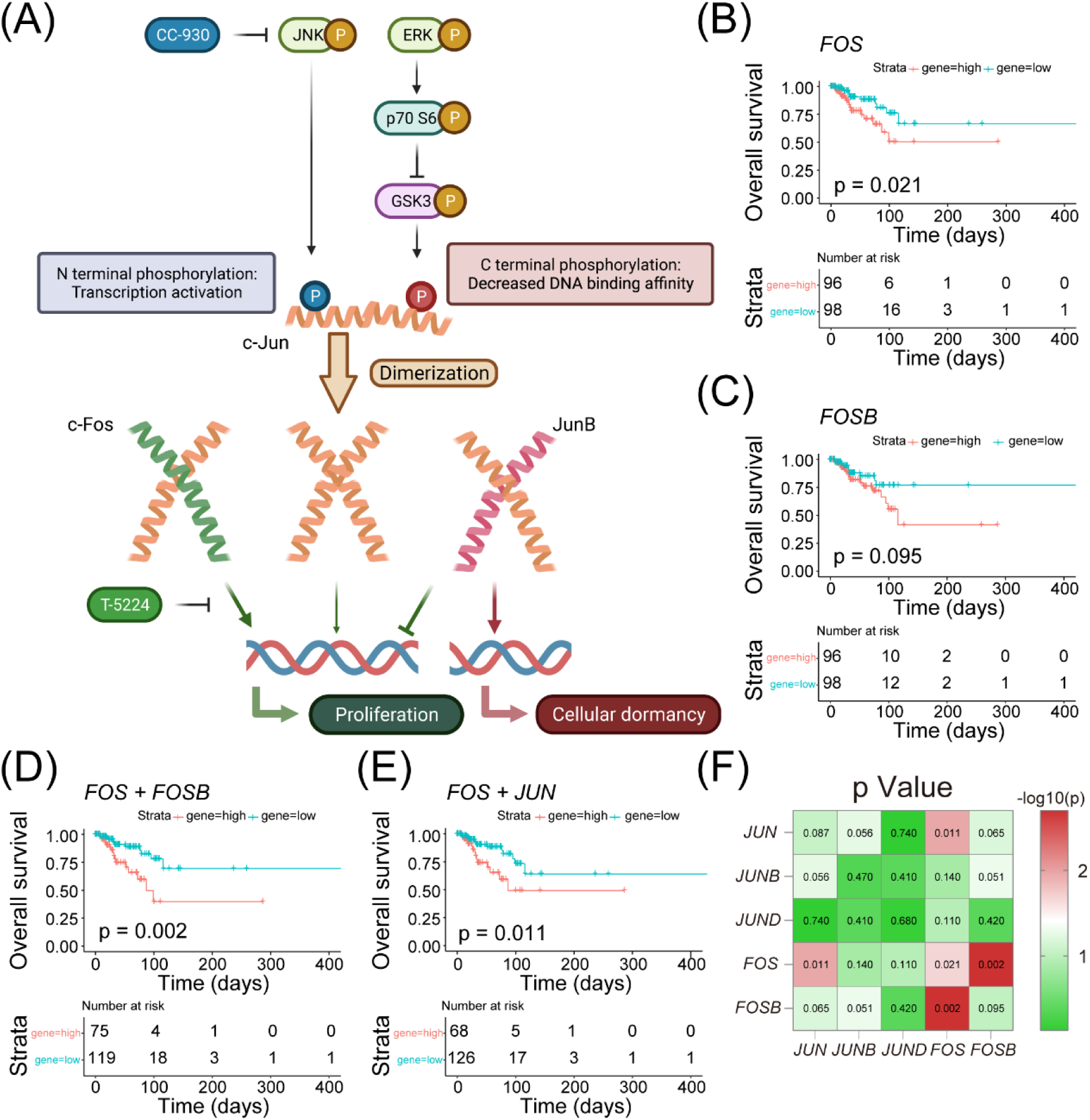
AP-1 overexpression was a poor prognostic factor in TNBC. (A) Schematic representation of AP-1 regulation in SCCs. c-Jun is at the center of AP-1 activity. N-terminal phosphorylation site of c-Jun contribute to AP-1 transcription activity, and C-terminal phosphorylation site of c-Jun prejudice to DNA binding affinity. Activation of JNK and ERK contribute to AP-1 transcription activity by phosphorylation of these two sites separately. c-Jun could form homodimer and form heterodimer dimer with c-Fos, promoting proliferation-related gene expression. JunB could from heterodimer dimer with c-Jun, inhibited proliferation-related gene expression through competitive binding, and promoting dormancy-related gene expression such as *CDKN1A* and *GADD45A*. (B, C) High expression level of *FOS* and *FOSB* was poor prognostic factor for TNBC patients. Patients with both gene expression > median were considered as high-expression group. Data was acquired from TCGA-BRCA database. n = 111 each, total n =222. (D) TNBC patient with both *FOS* and *FOSB* high expression had worse prognosis. Patients with both *FOS*/*FOSB* gene expression > median gene expression were considered as high-expression group. Data was acquired from TCGA-BRCA database. n=80, total n=222. (E) TNBC patient with both *FOS* and *JUN* high expression had worse prognosis. Patients with both *FOS*/*FOSB* gene expression > median gene expression were considered as high-expression group. Data was acquired from TCGA-BRCA database. n=74, total n=222. (F) The effect of simultaneous high expression of AP-1 gene on the survival of TNBC patients. Patients with two AP-1 subunits gene expression > median gene expression were considered as high-expression group. The data on the diagonal line is the effect of a single gene on patient survival. Data was acquired from TCGA-BRCA database, total n=222.

### 3.6 AP-1 regulate TNBC cellular dormancy and chemo-resistance

To delve deeper into the mechanism by which upregulation of AP-1 subunits affects cellular dormancy, we employed an in vitro chemo-treatment model to investigate whether inhibition of AP-1 could mitigate TNBC’s chemo-resistance induced by SCCs. We utilized a combination of docetaxel, doxorubicin, and cyclophosphamide, administered at clinically relevant dosages, which we henceforth refer to as TAC. Following 3 days of TAC treatment, the remaining cells were harvested. We observed an increase in all three subpopulations of SCCs after TAC treatment, with each population expanding from approximately 1% to 6.17 ± 0.44%, 50.63 ± 0.80%, and 8.88 ± 0.51%, respectively. Notably, the H2B-eGFP-enriched SCCs exhibited the most prominent increase, suggesting that doxorubicin played a significant role in the enrichment of SCCs. Subsequently, we assessed the efficacy of two AP-1-related small molecule inhibitors: CC-930, a selective c-Jun N-terminal kinase (JNK) inhibitor[41], and T-5224, a c-Fos/c-Jun DNA binding inhibitor[42]. Treating MDA-MB-231 cells with these inhibitors in combination with TAC, we found that CC-930 reduced the enrichment of SCCs resulting from TAC treatment, decreasing to 5.27 ± 0.27%, 18.13 ± 0.42%, and 7.74 ± 0.38%, respectively. Conversely, T-5224 did not decrease this enrichment; instead, it increased the populations of H2B-eGFP-enriched and Violet-enriched SCCs to 61.90 ± 0.46% and 10.01 ± 0.20% (Fig. 6A and Supplementary Fig. 8). Furthermore, by driving firefly luciferase expression under the AP-1 response element, we established a dual-luciferase assay to measure AP-1 transcriptional activity. After 24 hours of CC-930 treatment, relative firefly luciferase activity decreased, whereas cells treated with T-5224 showed no significant change compared to the control group (Supplementary Figure 9A). Taken together, these data suggest that CC-930 can mitigate the enrichment of SCCs during chemo-treatment by inhibiting AP-1 transcriptional activity.

**Fig 6:**
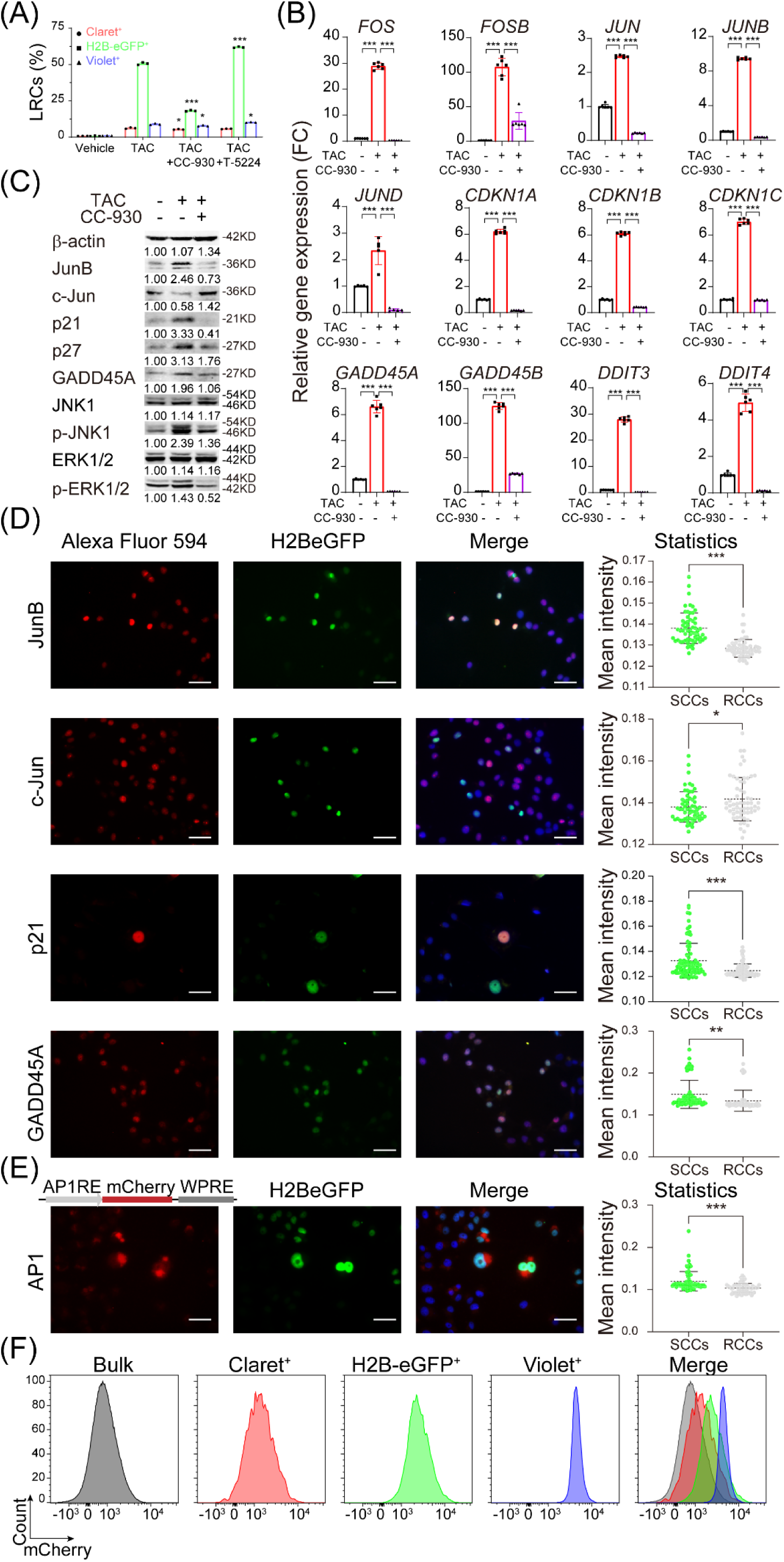
Mechanisms of AP-1-mediated anti-stress signaling in regulating TNBC SCC properties. (A) CC-930 could reduce SCCs residual caused by chemotherapy in vitro. TAC was a combination of docetaxel, doxorubicin, and cyclophosphamide, with common clinical dosage. Tri-labeled MDA-MB-231 cells were chased for 9 days, and TAC, CC-930 or T-5224 were added to each group. After 3 days of culture cells were collected and analyzed with flow cytometry. (B, C) CC-930 could reduce gene expression and protein level upregulated by TAC treatment. MDA-MB-231 cells were treated with TAC and 10 mM CC-930 for 3 days. Relative gene expressions were analyzed by RT-qPCR (B), and protein level were analyzed by Western Blot (C). (D) JunB, p21 and GADD45A were enriched in H2B-eGFP^+^ SCCs. Tet-On H2B-eGFP transfected MDA-MB-231 cells were incubated with doxycycline for 3 days and chasing without doxycycline for 12 days. Cells were fixed and permeabilized, and stained with corresponding antibody. Scale bar = 20 μm. (E, F) SCCs were enriched in AP-1 transcriptional activity. Tet-On H2B-eGFP and AP-1 reporter dual transfected MDA-MB-231 cells were incubated with doxycycline for 3 days and chasing without doxycycline for 12 days. (E) Cells were fixed and permeabilized, and stained with anti-mCherry antibody. Scale bar = 20 μm. (F) Equal amount of bulk cells, Claret^+^ SCCs, H2B-eGFP^+^ SCCs and Violet^+^ SCCs were collected and mCherry intensity were analyzed by flow cytometry. Data are represented as mean ± SD. *p ≤0.05; **p ≤0.01; ***p ≤0.001, Student’s t test.

We further investigated both gene and protein expression following TAC treatment. qPCR analysis revealed that *FOS* and *FOSB* exhibited the most significant upregulation after TAC treatment, with a 32.12-fold change and a 193.50-fold change, respectively. Members of the Jun family, including *JUN*, *JUNB*, and *JUND*, were also upregulated, with *JUNB* showing the highest increase at a 9.54-fold change. Additionally, we examined gene expression changes in the Cip/Kip family and GADD45s. Our findings indicated upregulation of genes associated with AP-1 subunits, the Cip/Kip family, and GADD45s after TAC treatment, which were subsequently reduced upon CC-930 treatment (Fig. 6B). Previous studies have indicated that c-Jun possesses at least two phosphorylation sites, located in both the N-terminal and C-terminal regions. N-terminal phosphorylation enhances c-Jun’s transcriptional activity and is mediated by JNK[43]. On the other hand, C-terminal phosphorylation reduces the binding affinity between the c-Jun dimer and DNA, and is catalyzed by GSK3 or CKII, with inhibition via cascade phosphorylation of ERK and p70-S6[44]. In short, both JNK phosphorylation and ERK phosphorylation both contribute to AP-1 transcriptional activity. Therefore, we also performed Western Blot detection on MDA-MB-231 cells treated with TAC or combined with CC-930. We found up-regulation in both JNK and ERK’s phosphorylation level after TAC treatment, and could be reversed by CC-930. It’s also worth noticing that, although *FOS* and *FOSB* gene expression had most obvious up regulation after TAC treatment, but there was no corresponding increase in protein level. Instead, the protein level of c-Fos, FosB were down regulated after TAC treatment, along with c-Jun (Fig. 6C). Considering the consistency of JunB at both RNA and protein levels, and the contradiction of c-Jun and c-Fos, we hypothesized that among multiple AP-1 subunits, JunB may be the most critical factor regulating dormancy.

To verify our speculation, we initially observed the co-expression of JunB and other related proteins in SCCs cells. We found co-localization between JunB expression, p21, GADD45A and SCCs marker H2B-eGFP (Fig. 6D). Also, we didn’t find much expression of c-Jun in SCCs, which parallels the findings observed in the previous Western blot results. We further developed an additional lentiviral AP-1 reporter system, wherein mCherry expression was driven by the AP-1 response element[45]. Stable MDA-MB-231 cells transfected with this construct were labeled with Claret, H2B-eGFP, and Violet as previously described, and then subjected to a 12-day chase period. Following the chase, the AP-1 transcriptional activity in each subpopulation of SCCs was assessed based on the intensity of mCherry fluorescence. We not only observed stronger AP-1 transcriptional activity in SCCs but also found variations in AP-1 transcriptional activity among different subpopulations of SCCs. Specifically, mCherry fluorescence intensity was most pronounced and specific in Violet-enriched SCCs, followed by H2B-eGFP-enriched SCCs and Claret-enriched SCCs (Fig. 6F). Immunofluorescence analysis further confirmed the co-localization between AP-1-derived mCherry expression and H2B-eGFP (Fig. 6E). Importantly, despite minimal differences in AP-1 expression levels, there was significant variability in AP-1 function among the three subpopulations of SCCs. Collectively, these findings underscore the involvement of AP-1 in the regulation of cellular dormancy in TNBC, suggesting that inhibition of AP-1 could effectively disrupt the dormant state of SCCs.

### 3.7 JUNB regulate TNBC cellular dormancy by directly activating the gene expression of CDKN1A and GADD45A

To comprehend the specific mechanism by which JunB regulates dormancy, we conducted subsequent experiments. Initially, a paired TMA experiment was performed to investigate JunB expression in patients with TNBC. Our findings revealed significantly higher JunB expression in the TNBC tumor group compared to the tumor adjacent group (Fig. 7A, B). In the subsequent steps, given the transcriptional activity of AP-1, we explored whether JunB could directly activate the expression of genes related to dormancy. We extracted promoter region sequences of the Cip/Kip family and GADD45s and analyzed them using the JASPAR database[46]. Our analysis indicated the presence of AP-1 binding sites in the promoter region of both the Cip/Kip family and GADD45s, with higher affinity observed on both the *CDKN1A* and *GADD45A* promoters (Fig. 7C and Supplementary Table S4). We then constructed another dual-luciferase assay by driving firefly luciferase expression under either the *CDKN1A* promoter or *GADD45A* promoter. We compared the relative firefly luciferase activity when transfected with either the JunB overexpression vector or the control vector. A higher relative firefly luciferase activity was observed when the JunB overexpression vector was transfected, indicating that high levels of JunB directly activated the transcription of either *CDKN1A* or *GADD45A* (Supplementary Fig. 9B). Furthermore, when the JunB overexpression vector was co-transfected with the previously constructed AP-1 response dual-luciferase system, we observed stronger relative firefly luciferase activity, indicating that overexpression of JunB could initiate AP-1 transcription (Supplementary Fig. 9C).

**Fig 7:**
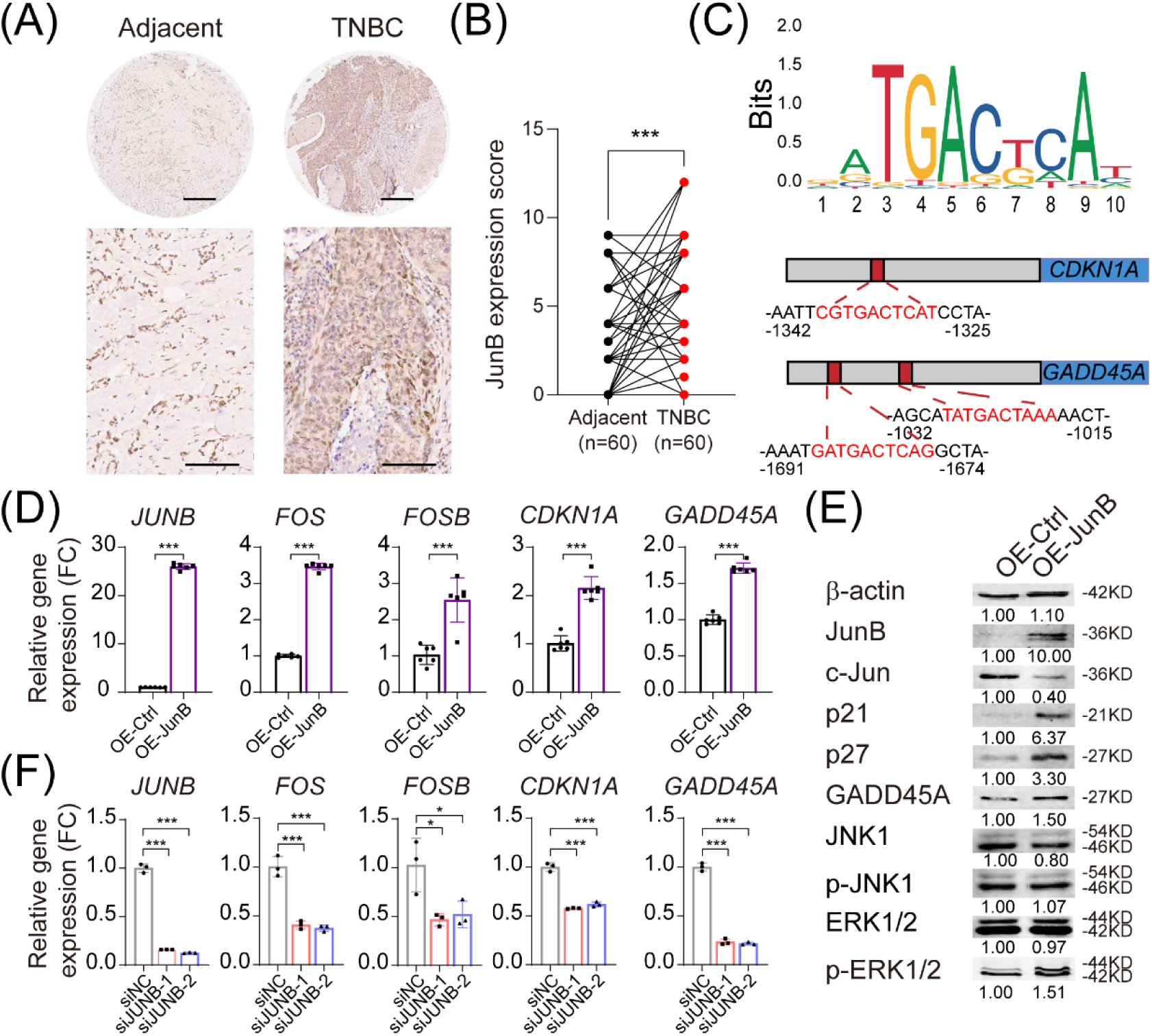
JunB was the key regulator of tumor dormancy. (A, B) JunB expression in TNBC patients. TMA with 60 paired tumor adjacent sample and TNBC sample was stained with anti-JunB antibody. JunB expression of each sample was scored and compared pairwise. Scale bar = 300 μm in the upper panel and 100 μm in the lower panel. (C) Schematic representation of predicted c-Jun/JunB heterodimer binding sites on CDKN1A and GADD45A promoter region. (D, E) Overexpression of JunB leaded to up regulation of Cip/Kip and GADD45s. Relative gene expressions were analyzed by RT-qPCR (D), and protein level were analyzed by Western Blot (E). (F) Interfere with JunB would result in down regulation of AP-1 subunits, Cip/Kip and GADD45s. Relative gene expressions were analyzed by RT-qPCR. Data are represented as mean ± SD. *p ≤0.05; **p ≤0.01; ***p ≤0.001, Student’s t test.

To confirm the regulatory role of JunB on CDKN1A and GADD45A, we established a stable MDA-MB-231 cell line overexpressing JunB. We observed upregulation of gene expression levels of *JUNB*, *FOS*, *FOSB*, *CDKN1A*, and *GADD45A* in JunB overexpression cells (Fig. 7D). Western blot analysis confirmed the upregulation of p21 and GADD45A in JunB overexpression cells, along with increased levels of phosphorylated ERK1/2 (Fig. 7E). Subsequently, we synthesized two siRNAs targeting JUNB and transfected them into MDA-MB-231 cells, observing an opposite gene expression pattern compared to JunB overexpression. Specifically, AP-1 subunits *FOS*, *FOSB*, and *JUN* were downregulated, while *CDKN1A*, *CDKN1B*, *GADD45A*, and *GADD45B* were also downregulated (Fig. 7F). These findings suggest that the c-Jun/JunB heterodimer can transcriptionally activate the Cip/Kip family and GADD45s. Taken together, all these data support the conclusion that upregulation of JunB in SCCs is a major factor contributing to cellular dormancy.

### 3.8 CC-930 could reduce SCCs residual after chemotherapy in vivo

To investigate the potential of anti-SCC therapy for clinical TNBC treatment, we evaluated CC-930’s anti-tumor efficacy both in vitro and in vivo. Previous experiments demonstrated that CC-930 reduced the residual SCCs after chemotherapy. In the colony formation assay, SCCs exhibited higher colony formation capacity than bulk cells, and CC-930 reduced colony formation (Supplementary Fig. 10C).

Further investigating CC-930’s anti-SCCs activity in vivo, we established 40 CDX models with MDA-MB-231 cells. Nude mice bearing CDX were randomly divided into four groups: control, CC-930 treatment, TAC treatment, or TAC + CC-930 combination for 21 days. Initially, tumors treated with TAC shrank but relapsed after 14 days. CC-930 treatment alone showed limited anti-tumor capacity but was effective when combined with TAC. The combination significantly reduced tumor size and weight, preventing tumor regrowth (Fig. 8A, B). As treatment progressed, mice in the TAC and combination groups exhibited a tendency of decreased body weight, which was not observed in the CC-930 treatment group (Supplementary Figure 10B). Abdominal metastases occurred in the control group (6 out of 10) and TAC group (6 out of 10), while no tumors in the CC-930 group developed abdominal metastases, with only 1 metastasis in the combination group (Fig. 8C). Finally, using SynergyFinder, we calculated that TAC and CC-930 exhibited a significant synergy effect (Fig. 8D). These findings suggest that CC-930 could enhance survival in TNBC patients treated with traditional chemotherapy.

**Fig 8:**
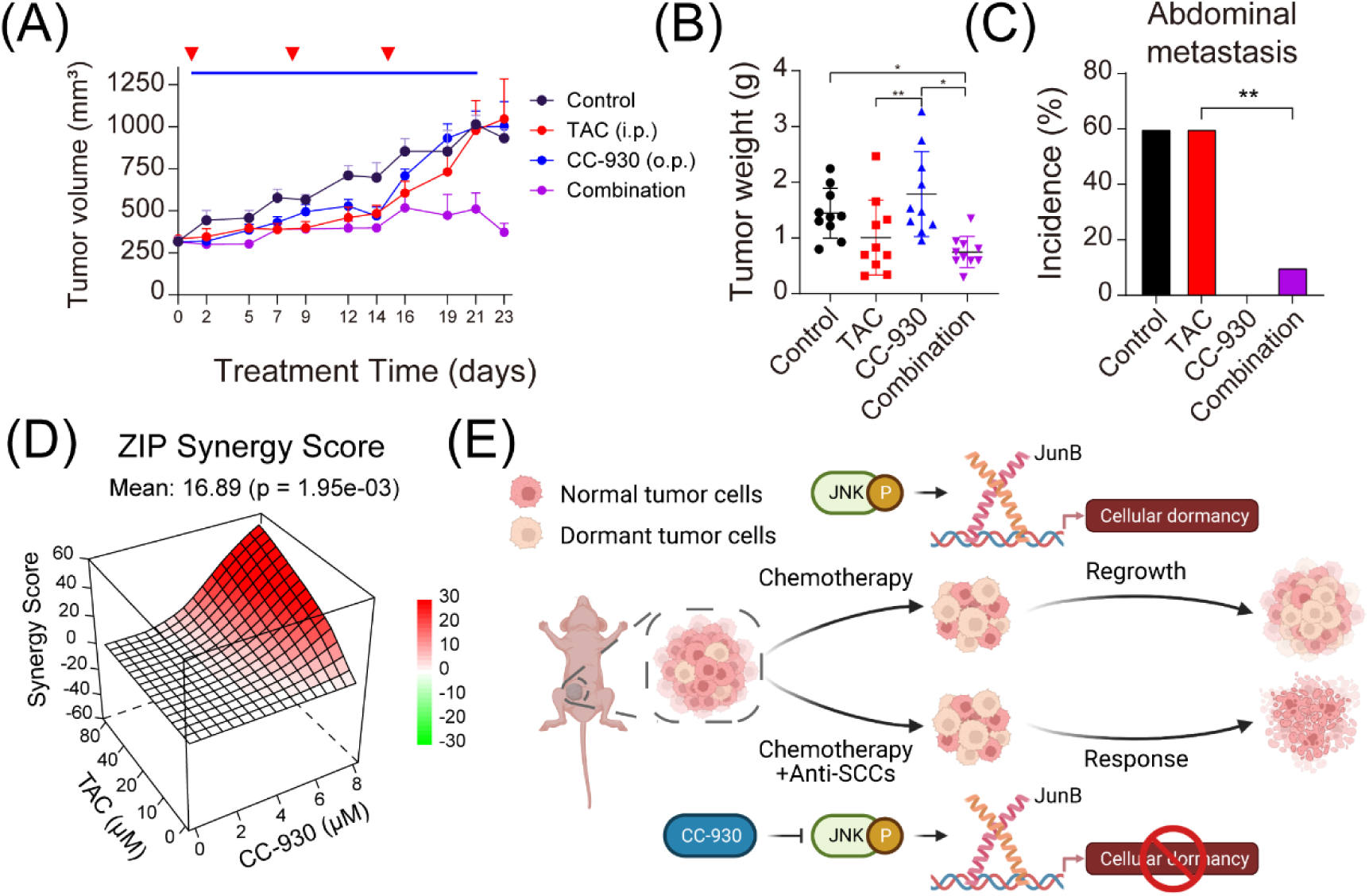
CC-930 could reduce SCCs residual after chemotherapy in vivo. (A, B) Timeline of in vivo experiment. MDA-MB-231 cells were injected into forth mammary fat pad in the right flank of nude mice. After 1 week, 40 mice were randomly divided into 4 groups, treated with vehicle, TAC, CC-930 or combination. TAC was given by i.p. every week, and CC-930 was given orally every day. Tumor length (L), tumor width (W) and body weight were measured 3 times a week. Tumor volume was calculated by V = (L x W x W)/2. After 21 days of treatment, mice were sacrificed, and tumors were removed and tumor weights were measured. CC-930 combined with TAC could effectively reduce the tumor volume (A) and tumor weight (B). (C) CC-930 could reduce abdominal metastasis combined with TAC. n = 10 in each group. (D) Synergy score of TAC and CC-930. Tumor volume data was annotated as viability data and were loaded into the free software Synergy Finder Plus. Synergy Score was calculated by ZIP synergy scoring. (E) Dormant TNBC cells were enriched in AP-1 activity and could evade chemotherapy. CC-930 could target the transcriptional activity of AP-1, resulting in more efficient clearance of TNBC.

## 4 DISCUSSION

Label retention is one of the most commonly used methods for labeling and enriching SCCs, allowing for the capture of live SCCs for subsequent experiments[22–29]. Different labeling methods employ various fluorescent dyes or fluorescent proteins, prompting a natural question: do different labeling methods yield distinct SCC subpopulations? Our chosen fluorescent labels include lipophilic dyes, protein covalent-binding dyes, and chromatin-tagged fluorescent proteins. These substances belong to the most common components in cells-phospholipid bilayers, chromatin, and proteins, respectively[47]. Firstly, we found that the same type of fluorescent labeling enriches highly correlated SCC subgroups. Various labeling retaining methods offer alternative fluorophores, which can be interchanged without additional impact when faced with limited fluorescent channel availability. More importantly, we found that different types of fluorescent labeling tended to enrich overlapping but not entirely identical SCCs. These SCCs exhibit some phenotypic similarities; for instance, they are all in a slow-cycling state and relatively insensitive to chemotherapy drugs, consistent with previous reports. More importantly, we discovered heterogeneity among these three groups of SCCs both at the distribution and functional levels. The SCCs subpopulation enriched with Claret and that enriched with Violet exhibit substantial differences, while the subpopulation enriched with H2B-eGFP falls between them. Violet enriched SCCs exhibit a more dormant phenotype, characterized by increased G1 phase arrest and enhanced drug resistance. In contrast, Claret enriched SCCs display more G2/M phase arrest, accompanied by weaker drug resistance but stronger migratory capabilities. One interpretation is that different subtypes of SCCs are in varying states of epithelial-mesenchymal transition (EMT)[48,49]. Another interpretation is associated with the enrichment of polyploid giant cancer cells (PGCCs) within SCCs.

PGCCs are a distinct subpopulation of cancer cells characterized by their abnormal DNA content, specifically having multiple sets of chromosomes (polyploidy), and unusually large cell size (giant morphology)[50]. These cells often arise from failed or aberrant cell division processes, such as endoreplication, where DNA replication occurs without subsequent cell division, leading to the accumulation of multiple copies of chromosomes within a single cell[51–53]. PGCCs have been observed in various types of cancers and are associated with tumor aggressiveness, resistance to therapy, and increased metastatic potential[53–56]. PGCCs typically exhibit a dormant phenotype, while their derived daughter cells demonstrate highly invasive and migratory characteristics[57]. We identified the presence of PGCCs within SCCs using PI staining (Fig 2D), and further confirmed the existence of PGCCs in SCCs through quantification of genomic DNA content and immunofluorescence analysis (Fig 6D, E). We found that Violet enriched SCCs exhibited the highest proportion of PGCCs, consistent with the abnormal protein accumulation observed in PGCCs. Meanwhile, certain characteristics of Claret enriched SCCs resembled those of daughter cells derived from PGCCs, particularly the enhanced migratory capacity and G2/M phase arrest. Hence, we speculate that the partial reason for the generation of distinct subtypes of SCCs lies in the differential distribution of various substances during the asymmetric division process of PGCCs. And also, the differences observed in drug resistance to chemotherapy among the three subpopulations, particularly the reactions of Violet enriched SCCs and H2B-eGFP enriched SCCs to taxanes and anthracyclines, can be explained by the theory. Taxanes could cause cytokinesis failure, and anthracyclines could cause karyokinesis failure, forming different forms of PGCCs[58,59]. Regarding the lack of enrichment of SCCs in response to PARP inhibitors and ATR inhibitors, it may be attributed to the p53 mutation in MDA-MB-231 cells[60,61], rendering the cells insensitive to these drugs and thus failing to induce significant cytotoxicity[62,63].

Another focus of this study lies in the analysis of the signaling pathways regulating dormancy in TNBC. Through RNA-seq, we found that SCCs exhibited downregulation in pathways related to cell cycle, DNA replication, and DNA damage repair, while showing upregulation in the MAPK and PI3K pathways, which is consistent with previous reports[64]. We also identified some expression characteristics of SCCs, such as ECM organization, angiogenesis, oxidative stress response, and protein misfolding, which are consistent with previously reported features of dormant cells[15,16,65–67]. It is noteworthy that we discovered that the function of AP-1, particularly the molecular function of JunB, is a key regulatory factor in cell dormancy.

The AP-1 is mainly composed of the c-Jun and c-Fos families. c-Jun, as the active center of AP-1, can form homodimers and initiate transcription. c-Fos can form heterodimers with c-Jun, and its transcriptional activity is higher than that of c-Jun homodimers. Both c-Jun homodimers and c-Fos/c-Jun heterodimers can promote cell proliferation. JunB and JunD can form heterodimers with c-Jun. The heterodimers formed have a higher DNA affinity compared to c-Fos/c-Jun dimers and can competitively counteract c-Jun-mediated cell proliferation[68–70]. The AP-1 transcription factor has been extensively studied, particularly its crucial role in regulating cell proliferation and survival[71,72]. Additionally, mounting evidence indicates that AP-1 is involved in multiple processes such as inflammation, differentiation, apoptosis, cell migration, and wound healing[68,71,73]. Recently, it has also been discovered that AP-1 supports angiogenesis, thereby enhancing tumors’ ability to cope with harsh microenvironments resulting from hypoxia[74,75]. Regarding cellular heterogeneity, recent evidence suggests that AP-1 also exerts a significant influence on the plasticity of tumor cells[76–91]. A most recent study in prostate cancer shows that FOXA2 collaborates with JUN on chromatin to promote transcriptional reprogramming of AP-1 in lineage plastic cancer cells, thereby facilitating transitions between multiple lineages of cellular states[86]. Our research team also investigated the connection between AP-1 and FOX family members in prostate cancer through single-cell chromatin landscapes, identifying them as critical regulatory factors in castration response[80].

Furthermore, we found that c-Jun/JunB heterodimers can directly activate the transcription of genes such as *CDKN1A* and *GADD45A*, thereby blocking the cell cycle. It is worth noting that we observed both higher RNA and protein levels of JunB in SCCs, while there were discordant RNA and protein levels of c-Jun and c-Fos. We speculate that this is because JunB competes with c-Jun for binding, leading to rapid degradation of c-Fos and FosB, as c-Fos is a highly unstable protein with rapid nuclear-cytoplasmic shuttling[92]. In addition, recent research has shown that at least one of the three known c-Jun E3 ubiquitin ligases, Itch, is activated by JNK phosphorylation[93,94]. Elevated levels of Fos and FosB gene expression favor cell re-entry into the cell cycle and proliferation. When the external environment changes, high levels of gene expression can be efficiently translated into proteins, further promoting cell proliferation. These findings suggest that AP-1, especially JunB subunit, is involved in the regulation of TNBC dormancy. It is also noteworthy that this transition process may not only rely on JunB expression. He et al. reported that JunB can form a complex with YAP/TAZ, and JunB has fewer overlapping motifs with the traditional YAP/TAZ downstream TEAD[95]. This is consistent with previous reports that COL17A1 maintains colon cancer stem cell dormancy by inhibiting YAP/TAZ[96]. In addition, AP-1 can also form complex dimers with other bZIP family members, such as the C/EBP family[69,97]. We also found some of the C/EBP family were highly expressed in SCCs (data not shown). These pieces of evidence indicate that the mechanism by which AP-1 regulates dormancy could be more complex than we described.

Finally, we discovered that CC-930 sensitizes SCCs to chemotherapeutic drugs by inhibiting AP-1 activity. When used alone, CC-930 did not exhibit significant tumor suppressive effects. However, it effectively prevented abdominal metastases and reduced abdominal metastases induced by TAC treatment in combination groups. It is noteworthy that CC-930 is a safe, orally active drug[41]. The dosage of CC-930 used in our study was relatively low compared to other in vivo assays published previously[98–100]. It is possible that a higher dose of CC-930 could yield better results. These findings underscore the potential of anti-SCCs therapy to enhance survival in TNBC patients and highlight CC-930 as an effective treatment option for SCCs.

Our study also has several limitations. Firstly, our research focuses primarily on in vitro experiments using MD-MB-231 cells, with fewer in vivo experiments conducted. Indeed, many features observed in SCCs, such as elevated ECM expression and enrichment of aging-associated phenotypes, suggest their potential involvement in altering the tumor immune microenvironment. Our previous work has also confirmed the association of dormancy-related ECM with tumor immune suppression microenvironments[65]. Secondly, we lack more clinically relevant models, such as PDX or organoids. Lastly, the minimal gene expression differences among the three subtypes of SCCs could potentially be further elucidated by integrating data such as ATAC-seq to describe the differentiation processes of SCC subtypes.

## 5 CONCLUSIONS

In conclusion, we discovered that different label-retaining methods can label and enrich overlapping but not identical subpopulations of SCCs. These SCCs subpopulations are all cell cycle arrested, but they exhibit heterogeneity in cell cycle distribution, drug resistance, invasive and other aspects. We found that the high expression of AP-1 subunits is a poor prognostic factor specific to TNBC, and the AP-1 subunit JunB plays a crucial role in regulating TNBC dormancy. Intervention of AP-1 function with CC-930 in combination with chemotherapy drugs can reduce tumor mass and decrease abdominal metastasis, showing promising clinical applications.

## AUTHOR CONTRIBUTIONS

Dean G. Tang, Jianjun Zhou and Yang Dong designed this study. Yang Dong, Jin Bai, Anmbreen Jamroze, Rong Fu, Huilan Su, Shan Wu and Wenwen Xia performed the experiments. Dean G. Tang and Jianjun Zhou supervised the direction of project and interpreted the data. Dean G. Tang and Yang Dong wrote the manuscript. All authors discussed the results and commented on the manuscript.

## ACKNOWLEDGEMENTS

We thank Dr. Zuoren Yu, Dr Yandong Li and Dr Ke Wei (Shanghai East Hospital, Tongji University) and anonymous reviewers for reading and commenting on the manuscript.

## CONFLICT OF INTERESTS STATEMENT

The authors declare that they have no competing interests.

## FUNDING INFORMATION

This study was supported by the Chinese Ministry of Science and Technology (MOST) Grant (2016YFA0101201)

## ETHICS APPROVAL

The research on the use of human tissue samples was approved by the Medical Ethics Committees of Shanghai East Hospital, Tongji University.

## CONSENT FOR PUBLICATION

Not applicable.

## DATA AVAILABILITY STATEMENT

The RNA-seq data have been deposited in GEO database under accession code GSE267757 and GSE267759

## Notes

### Competing Interest Statement

The authors have declared no competing interest.

### Summary of Updates

Author list correctionsAuthor list corrections；Draft updates

